# PL1 Peptide Engages Acidic Surfaces on Tumor-Associated Fibronectin and Tenascin Isoforms to Trigger Cellular Uptake

**DOI:** 10.1101/2021.09.16.460582

**Authors:** Prakash Lingasamy, Kristina Põšnograjeva, Sergei Kopanchuk, Allan Tobi, Ago Rinken, Ignacio J. General, Eliana K. Asciutto, Tambet Teesalu

**Affiliations:** Laboratory of Precision and Nanomedicine, Institute of Biomedicine and Translational Medicine, University of Tartu, Tartu, 50411 Tartu, Estonia; Center for Nanomedicine and Department of Cell, Molecular and Developmental Biology, University of California, Santa Barbara, Santa Barbara, 93106 California, USA; Institute of Chemistry, University of Tartu, Ravila 14, Tartu, 50411, Estonia; School of Science and Technology, National University of San Martin (UNSAM), ICIFI and CONICET, 25 de Mayo y Francia, 1650 San Martín, Buenos Aires, Argentina

**Keywords:** PL1 peptide, fibronectin extra domain B (FN-EDB), Tenascin-C C domain (TNC-C), molecular docking, molecular dynamics simulations, dissociation constant, glioblastoma, extracellular matrix (ECM), phage display, nanoparticles

## Abstract

Tumor extracellular matrix (ECM) is a high-capacity and genetically stable target for the precision delivery of affinity ligand-guided drugs and imaging agents. Recently, we developed a PL1 peptide (sequence: PPRRGLIKLKTS) for systemic targeting of malignant ECM. Here we map the dynamics of PL1 binding to its receptors Fibronectin Extra Domain B (FN-EDB) and Tenascin C C-isoform (TNC-C) by computational modeling and cell-free binding studies on mutated receptor proteins, and study cellular binding and internalization of PL1 nanoparticles in cultured cells. Molecular dynamics simulation and docking analysis suggested that the engagement of PL1 peptide with both receptors is primarily driven by electrostatic interactions. Substituting acidic amino acid residues with neutral amino acids at predicted PL1 binding sites in FN-EDB (D52N-D49N-D12N) and TNC-C (D39N-D45N) resulted in the loss of binding of PL1 nanoparticles. Remarkably, PL1-functionalized nanoparticles (NPs) were not only deposited on the target ECM but bound the cells and initiated a robust cellular uptake via a pathway resembling macropinocytosis. Our studies establish the mode of engagement of the PL1 peptide with its receptors and suggest applications for intracellular delivery of nanoscale payloads. The outcomes of this work can be used for the development of PL1-derived peptides with improved stability, affinity and specificity for precision targeting of the tumor ECM and malignant cells.

**One Sentence Summary:** PL1 peptide is recruited to the acidic surfaces on oncofetal fibronectin EDB and tenascin C-C isoform, triggering cellular uptake of PL1-functionalized nanoparticles.

## Introduction

Systemic homing peptides are used for the precision delivery of drugs and nanoparticles (NPs). In addition to affinity targeting of plasma membrane receptors expressed on the surface of malignant and tumor-associated cells, homing peptides with specificity towards tumor-associated ECM molecules have been developed [1–6]. In tumors, ECM is an abundant high-capacity target that is primarily deposited by genetically stable stromal cells and not by rapidly mutating malignant cells. ECM is a dynamic structure and its composition reflects changes in the tissue status and disease state. For example, fibronectin has over 20 different alternatively spliced isoforms, and some of these variants are expressed only during embryonic development and in activated tissues, including solid tumors. The expression of extra domain B of fibronectin (FN-EDB) and tenascin-C C isoform (TNC-C) is associated with angiogenesis and tissue remodeling, whereas resting normal adult tissues are negative [2–4,7,8]. FN-EDB and TNC-C are among the most specific markers for angiogenic blood vessels [9]. In tumors, the malignancy and poor prognosis correlate with FN-EDB and TNC-C expression [10–12]. Monospecific affinity targeting ligands have been reported for both FN-EDB [e.g., FN-EDB-ScFV L19 [13], APT_EDB_ [14], ZD2 [15]], and TNC [e.g., TNC-C-ScFV G11 [2,4], TNC aptamer [16], and TNC-binding FHK peptide [17]]. Some of these targeting ligands are undergoing clinical testing for precision imaging and therapy of solid tumors [18–21].

Recently, we developed a bispecific PL1 peptide (amino acid sequence PRRGLIKLKTS) that recognizes both FN-EDB and TNC-C [3]. Compared to monospecific targeting, simultaneous targeting of FN-EDB and TNC-C has the advantages of increased binding capacity and more uniform tumor delivery. In preclinical studies, the intravenously-injected PL1 nanoparticles showed robust accumulation in a panel of glioblastoma and prostate carcinoma xenografts models in mice, and the treatment of glioblastoma mice with proapoptotic PL1 nanoparticles resulted in suppression of tumor growth and extended survival [3,21].

We used molecular dynamics to study the interactions that drive the binding of PL1 to its receptors by constructing molecular models. These studies revealed a preponderance of electrostatic interactions, which was also confirmed experimentally. Finally, we evaluated the suitability of the peptide for intracellular cargo delivery. Our study provides a basis for follow-up translational work to enhance the stability, specificity and affinity of the bioactive conformation of the PL1 peptide for improved affinity targeting of the tumor ECM and malignant cells.

## Materials and Methods

### Materials

Dulbecco’s Phosphate Buffer Saline (PBS; cat #D8662 or Lonza, cat #BE17-516F), Pluronic F-127 (cat #P2443) and isopropyl β-D-1-thiogalactopyranoside (IPTG; Sigma, cat #I6758) were purchased from Sigma-Aldrich (Munich, Germany). The PL1 peptide (PPRRGLIKLKTS) with free carboxyl terminus and 5(6)-carboxyfluorescein (FAM) or biotin coupled via the 6-aminohexanoic acid spacer to the N-terminus of the peptide was ordered from TAG Copenhagen (Frederiksberg, Denmark). The peptide nanoparticles, iron oxide nanoworms (NWs) and silver nanoparticles (AgNPs), were prepared as described previously [3,8,22,23].

### Computational modeling and molecular dynamics simulations

The NMR structure (PDB 2FNB) of the human oncofetal fibronectin EDB [24] was used to build the initial FN-EDB model. In the case of TNC-C, there was no high-resolution structure available and a model was built using the I-TASSER online server [25] for protein structure prediction. Electrostatic analyses of these models were performed using the APBS web-server [26].

Molecular dynamics simulations of the aforementioned and mutated models (where selected charged residues were mutated, as detailed in the Results section) were done using the AMBER18 [27] software package, with all of them—except for the minimizations—using the GPU version of the PMEMD program, and the Amber 14 force field [28] was employed. The protocol used for minimization and equilibration was: (1) 5,000 cycles of minimization using the steepest descent method, followed by another 5,000 steps using conjugate gradient; (2) 100 ps of heating to 300 K, using a Langevin thermostat with a collision frequency of 1.5 ps^−1^; (3) 5 ns with constant T (300 K) and P (1 atm), using a weak-coupling Berendsen barostat with a relaxation time of 2 ps; (4) A final equilibration in the NVT ensemble (constant volume and temperature) of 5 ns. The SHAKE algorithm was adopted in all simulations, allowing the use of a 2 fs time step, and long-range electrostatics were taken into account using periodic boundary conditions, via the Particle Mesh Ewald algorithm [29], with a cut-off of the sums in direct space of 12 Å. This protocol was used for all simulated systems for both FN-EDB and TNC-C. The succeeding production runs were done in the NVT ensemble with 1 μs duration.

### Recombinant protein expression and purification

Wild type (WT) TNC-C and FN-EDB were prepared as described previously [2,3,8]. The DNA inserts encoding for hexahistidine-tagged mutant TNC-C variants (TNC-C E48Q, TNC-C D39N D45N, TNC-C D13N E70Q E88Q) and mutant FN-EDBs variants (FnEDB D68N D70N, FnEDB D52N D49N D12N, FnEDB E42Q D68N E64Q) were synthesized and cloned into pET28a+ expression vectors (GenScript Biotech BV, Netherlands). The resulting plasmids were transformed into the *E. coli* BL21 Rosetta 2 (DE3) pLysS strain (Novagen, #70956). The protein expression was induced with 0.5 mM IPTG and followed by culturing at 20 °C for 16 h. The bacterial cells were lysed by sonication (Bandelin Sonopuls HD 2070, Merck, Germany) in ice cold IMAC buffer (25 mM Tris-HCL, 400 mM NaCl, 25 mM imidazole pH 8, protease inhibitors cocktail, DNase I, 1 mM MgCl_2_). The soluble fraction of the cell lysate was purified on HiTrap IMAC HP columns (GE Healthcare # 17-0920-05) according to the manufacturer’s protocol. The purified proteins were dialyzed against PBS using 3.5 kDa cut-off Slide-A-Lyzer Dialysis Cassettes (Thermo Scientific #66330). The protein concentration was measured by bicinchoninic acid (BCA) assay (Thermo Scientific #23227). The protein purity and size were analyzed using 15% SDS-PAGE.

### PL1 phage binding assay

For cell-free binding, His-6X-tagged proteins (30 μg/10 μl beads) in 400 μl of PBS were incubated with Ni-NTA Magnetic Agarose Beads (QIAGEN, Hilden, Germany) at room temperature (RT) for 1 h. The beads were washed 3 times with washing buffer (PBS, 1% BSA, 0.1% NP40), followed by incubation with T7 phage displaying PL1 and control peptides at a final concentration of 5 x 10^8^ pfu in 100 μl in washing buffer at RT for 1 h. The background phages were removed by rinsing 6 times with washing buffer, and the bound phage-receptor complexes were released with 1 ml of elution buffer (PBS, 500 mM Imidazole, 0.1% NP40). The eluted phages were titered for plaque-forming unit (PFU) counting. The phage clones displaying heptaglycine peptide (G7), or insertless phage were used as negative controls.

### Binding of PL1 nanoworms to recombinant proteins

The recombinant hexahistidine-tagged TNC-C and EDB WT and mutant proteins (at 60 μg of protein / 40 μl beads in PBS) were immobilized to Ni-NTA Magnetic Agarose Beads (QIAGEN, Hilden, Germany) at RT for 1 h. The protein-coated beads were washed 3 times with washing buffer (PBS, 1% BSA, 0.1% NP40). Fluorescently-labeled PL1-NW (80 μg of NWs/reaction) were incubated with protein-coated beads in the washing buffer. The free PL1-NWs were removed by rinsing 6 times with washing buffer. The protein-PL1-NWs complex was eluted with 1 ml of elution buffer (PBS, 500 mM Imidazole, 0.1% NP40). The fluorescence of eluted samples was quantified using a Victor X5 Multilabel Plate Reader (PerkinElmer, Massachusetts, USA) at an excitation wavelength of 485 nm and an emission wavelength of 535 nm.

### Assessment of binding affinity by fluorescence anisotropy in a multiwell microplate

Fluorescence anisotropy (FA) saturation binding experiments were performed as previously described [30,31]. The experiments were carried out in PBS with the addition of 0.1% Pluronic F-127 (Sigma-Aldrich, Cat#P2443) in a final volume of 100 μl using a 96-well half area flat-bottom polystyrene NBS multiwell plates (Corning, cat #3686). The different concentrations of proteins (0–112 μM FN-EDB or 0–275 μM TNC-C) were added to a fixed concentration (0.66 μM) of FAM-PL1. The total and non-specific binding was measured in the absence or in the presence of a 500 μM Biotin-Ahx-PL1, respectively, after 24-h incubation at 25 °C in the dark sealed with a moisture barrier (4Titude, Cat# 4ti-0516/96). The concentration of fluorescent ligand and proteins in stock solutions was determined by absorbance (for FAM-PL1 ε_495_ = 75000 M^−1^ cm^−1^, for FN-EDB ε_280_ = 11460 M^−1^ cm^−1^ and TNC-C ε_280_ = 8480 M^−1^ cm^−1^ were used). The measurements were performed at 25 °C on a Synergy NEO (BioTek) microplate reader using an optical module with an excitation filter at 485 nm (slit 20 nm), emission filter at 528 nm (slit 20 nm), and polarizing beam splitting for dual-channel detection. Dual emission detection mode allows simultaneous recording of parallel (I||) intensities and perpendicular (I⊥) to the plane of excitation light. Sensitivities of channels (G factor) were calibrated with gain adjustment of the photomultiplier tubes using fluorescein (1 μM reference solution, Lambert Instruments) as a standard. The fluorescence anisotropy values were calculated as parameters FA from the equation FA= (I_||_ - G·I⊥)/(I_||_ + 2·I⊥). The binding affinity was estimated by global fitting of the data as in [32]. This simultaneous fitting of total and non-specific binding data considers the ligand depletion by both binding processes.

### Cellular binding and uptake of PL1-AgNPs in glioma cells

Biotinylated PL1 peptide or only biotin as a non-targeted control and CF555 dye were conjugated to neutravidin-AgNPs. The U87-MG cells were cultured on glass coverslips until 80–90% confluency and incubated with PL1-AgNPs or control biotin-blocked AgNPs at 37 °C for 20, 60 and 180 min. The free AgNPs were removed by washing with culture medium. The cell surface-bound AgNPs were removed with a freshly prepared etching solution containing 10 mM of Na2S2O3 and K3Fe (CN)6 in PBS, which was applied for 3 mins. The cells were fixed with −20 °C methanol for 2 min and washed with PBS 3 times. In some of the samples, the cells were stained to visualize the target receptors FN-EDB and TNC-C with respective primary and secondary antibodies. The nuclei were counterstained with 4’,6-diamidino-2-phenylindole (DAPI, Molecular Probes) at 1 μg/ml. Fluoromount-G (Electron Microscopy Sciences, PA, USA) medium was used to mount the coverslips on microscope slides for confocal imaging (Zeiss LSM 510; Olympus FV1200MPE, Germany).

In addition, flow cytometry was performed to assess the binding and uptake of PL1-AgNPs. Attached U87-MG cells were incubated with PL1-AgNPs and control AgNPs in complete cell culture medium at 37 °C for 1 h. The cells were washed, and etched to remove noninternalized particles. The cells were analyzed using the BD Accuri flow cytometer by monitoring the 555 nm channel FL2.

For characterization of the AgNP uptake pathway in glioma cells, the U87-MG cells were cultured until 80–90% confluency and pre-incubated with the following endocytosis inhibitors: (1) 50 μM Nystatin to block caveolae/lipid-mediated endocytosis, (2) 50 μM 5-(N-ethyl-N-isopropyl)-Amiloride (EIPA) to block micropinocytosis, (3) 4 μM Cytochalasin D to block actin polymerization and micropinocytosis, and (4) 10 μM Chlorpromazine to block clathrin-mediated endocytosis. The PL1-AgNPs or control biotin-AgNPs were then added, and the cells were incubated at 37 °C for 1 h and subsequently washed to remove free particles. Some samples were also treated with an etching solution to remove accessible surface-bound AgNPs. The fluorescence of the labeled AgNPs was detected and analyzed using the BD Accuri 6 flow cytometer in the 555 nm channel.

### Statistical Analysis

Prism 6 software was used for statistical analysis. The results were presented as mean with error bars indicating ± SEM. For a comparison of 2 groups, we used an unpaired t-test, one-way ANOVA test, and ANOVA test. P < 0.05 was considered significant and P-values were indicted as follows: * - P ≤0.05, ** - P ≤0.01, *** - P ≤0.001 and **** - P ≤0.0001.

## 2. Results

### 2.1. PL1 binding to FN-EDB and TNC-C is electrostatically driven

To map the FN-EDB region(s) involved in PL1 binding, we first performed a docking analysis using the pepATTRACT software [33]. Candidate regions involved in peptide binding included the one with the largest affinity score, where PL1 was positioned head-on to D12 of FN-EDB. Next, we perturbed the peptide by moving it a few angstroms away from the docked position to see if the peptide returns to its original position, thus suggesting the stability of that conformation. We observed that following perturbation, PL1 did not go back to the initial position in 500 ns but docked not far from the initial binding site. In parallel, we performed eight independent molecular dynamics simulations with FN-EDB and a peptide located randomly in a water box far from each other. These simulations, ran for 2 μs each, showed that in all cases PL1 associated with the FN-EDB in no longer than 100 ns. Three specific regions of FN-EDB had higher affinity for the PL1 peptide than the rest of the molecule: (1) D12, D49 and D52; (2) D68 and D70; and (3) E42, E64 and D68 (Fig. 1A); these regions were also found via the docking analysis. The same approach was applied to model interactions of PL1 with TNC-C. For TNC-C, the PL1 peptide appeared to engage preferentially with the lower half of TNC-C with the involvement of the following TNC-C residues: (1) D13, E70 and E88; (2) D39 and D45; (3) E48 (Fig. 1B). As in the case of FN-EDB, these regions were also compatible with the ones found via docking.

**Figure 1.**
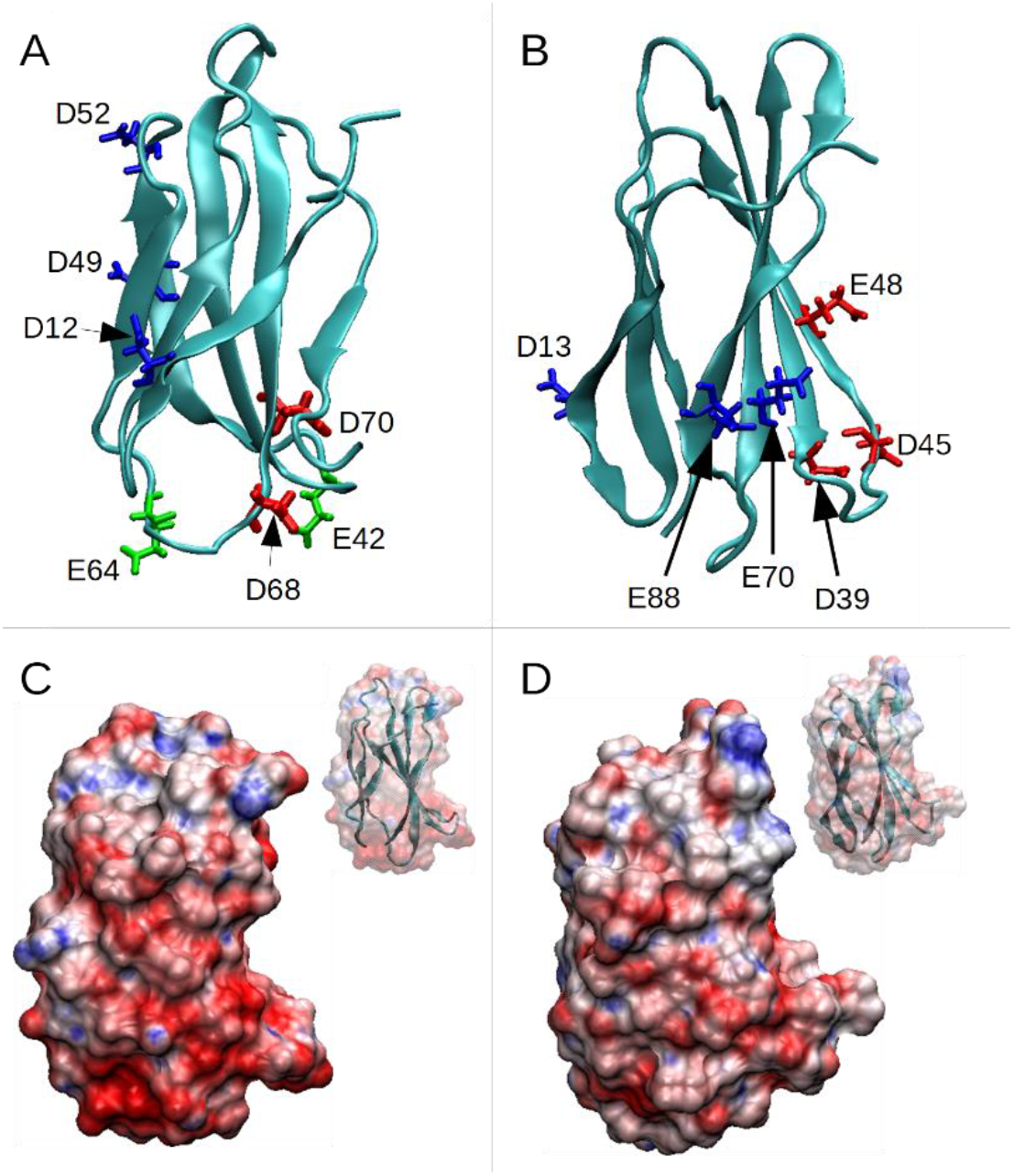
High-affinity residues and electrostatic potential of PL1 binding partners. (A) Structure of FN-EDB showing three areas of charged residues, (1) D52, D49 and D12; (2) D68 and D70; and (3) E42, D68 and E64. (B) Structure of TNC-C showing three areas of charged residues, (1) D13, E70 and E88; (2) D39 and D45; (3) E48. (C) Electrostatic potential surface of FN-EDB (red represents regions with a negative potential, while blue represents positive ones); the inset shows the cartoon representation of the protein. (D) Electrostatic potential surface of TNC-C.

Both FN-EDB and TNC-C have a net negative charge, of −10e and −5e (where e is the elementary charge), respectively. On the other hand, PL1 peptide has a net charge of +4e. Figures 1C, 1D display the electrostatic potential surface of the two proteins, with lower halves presenting a highly negative charge (in red) more prone to attracting the basic PL1 peptide.

To estimate the affinity of the PL1 peptide to the regions of FN-EDB and TNC-C established above, we took snapshots of the PL1-receptor complexes found above, and performed new molecular dynamics simulations using a *pulling procedure* [34], with a harmonic restraint (spring constant of 0.1 kcal·mol^−1^·Å^−2^) used to force the unbinding of the two molecules. This pulling was applied to three systems in FN-EDB:

i. Wild-type (WT)
ii. Mutations D12N, D49N, D52N (Set 1)
iii. Mutations D12N, D49N, D52N and D68N, D70N (Set 2)

and three systems in TNC-C:

i. Wild-type (WT)
ii. Mutations D13N, E70Q, E88Q (Set 1)
iii. Mutations D13N, E70Q, E88Q and D39N, D45N (Set 2)

Simulations were run for 20 ns, driving the systems from bound to unbound conformations. In each simulation the procedure was repeated 8 times to allow the system to unbind following different paths. Figures 2A and 2B show the results for the 3 systems (WT, Set 1 and Set 2), for FN-EDB and TNC-C, respectively.

**Figure 2.**
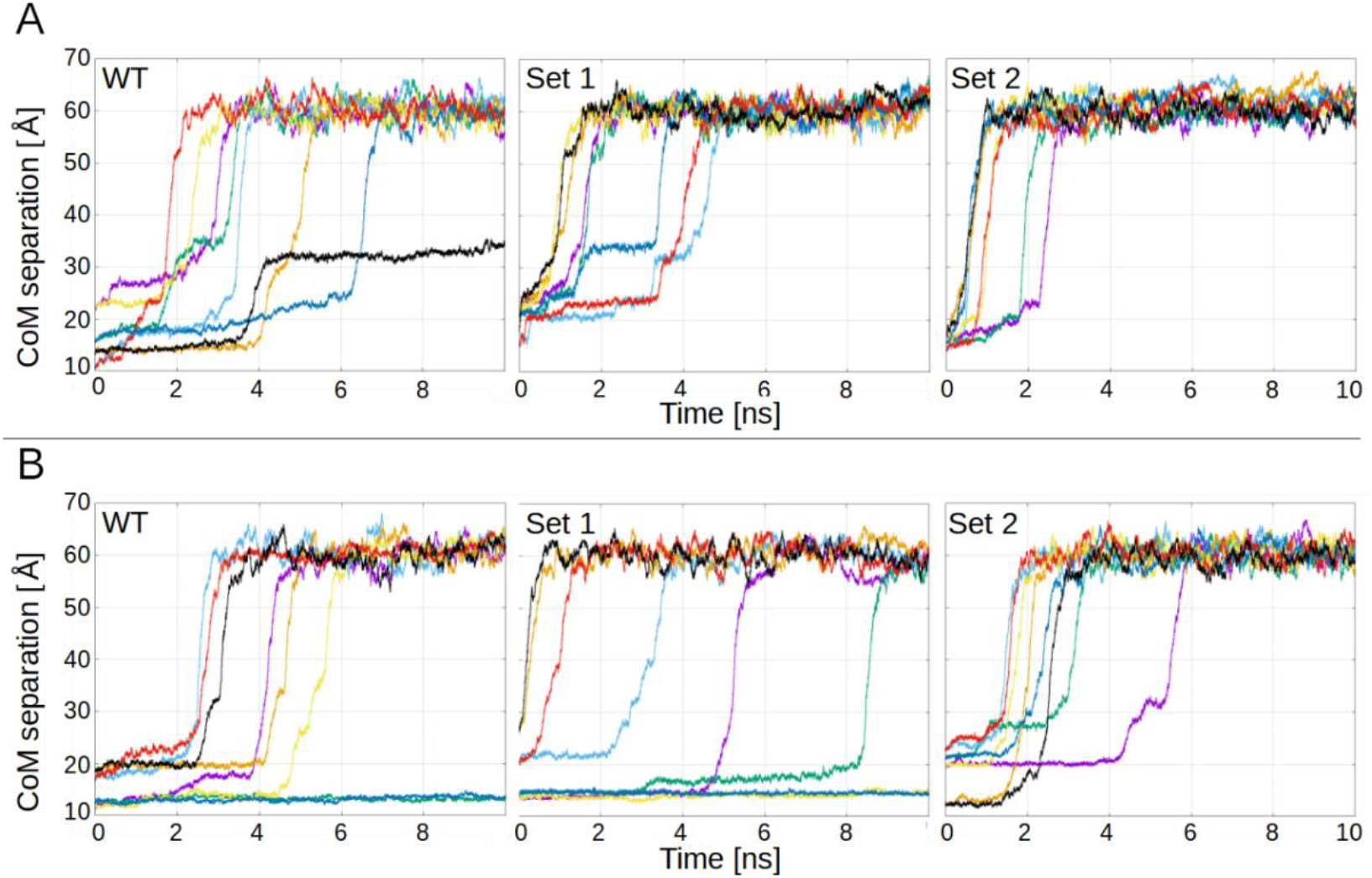
PL1 unbinding from receptors. (A) Harmonic restraint-driven unbinding of PL1 from WT FN-EDB, mutant Set 1 FN-EDB (D12N, D49N, D52N), and mutant Set 2 FN-EDB (D12N, D49N, D52N and D68N, D70N). (B) Harmonic restraint-driven unbinding of PL1 from WT TNC-C, mutant Set 1 TNC-C (D13N, E70Q, E88Q), and mutant Set 2 TNC-C (D13N, E70Q, E88Q and D39N, D45N).

Figure 2A shows the separation between the center-of-mass (CoM) of FN-EDB and PL1, as a function of simulation time. Each curve represents a different starting pose. Comparing the panels, the unbinding is stronger (occurs faster) in Set 2, takes more time in Set 1, and even more time in the WT. This shows that whereas the mutations in Set 1 favor unbinding (as compared to the WT), the mutations of Set 2 (which includes Set 1) favor it even more.

CoM separations of TNC-C and PL1 show a similar pattern: stronger binding in WT, followed by Set 1, and Set 2 being the weakest (Fig. 2B). In the case of WT TNC-C, there are two curves that do not show unbinding, and the rest do show it during the second quarter of the simulation time. Set 1 still shows two curves with no unbinding, one (green) that appears strongly bound but unbinds towards the end, and all the rest are weaker than in the case of WT, with three unbinding events in the first quarter. And finally, in Set 2, all systems but one show unbinding by the end of the first quarter. On average, the sequence of strong to weak binding is WT, Set 1 and Set 2, with Set 2 assuring full unbinding.

### 2.2. Under cell-free conditions, PL1 peptide engages acidic binding sites on TNC-C and FN-EDB

PL1 engages with its receptors TNC-C and FN-EDB in cell-free, *in vitro* and *in vivo* conditions [3,21,35]. The alanine scanning mutagenesis of PL1 peptide indicated the critical role of positively charged amino acids for TNC-C and FN-EDB binding [3]. These experiments as well as molecular docking and simulation studies described in the previous section highlight the potential roles of negatively charged binding surfaces on both FN-EDB and TNC-C for the engagement of PL1 peptide.

To test the *in silico* predictions, we tested PL1 binding to mutant TNC-C and FN-EDB in which the negatively charged amino acids in the predicted binding sites were substituted with neutral amino acids. The hexahistidine-tagged mutant FN-EDB (D68N D70N, D52N D49N D12N, and E42Q D68N E64Q) and TNC-C (E48Q, D39N D45N, and D13N E70Q E88Q) (Fig. S1) isoforms were expressed in *E. coli* and purified using affinity chromatography on Ni-NTA matrix (Fig. 3A). The purified recombinant proteins were coated on Ni-NTA magnetic beads, and the binding of PL1 and control nanoparticles was quantitatively studied.

**Figure 3.**
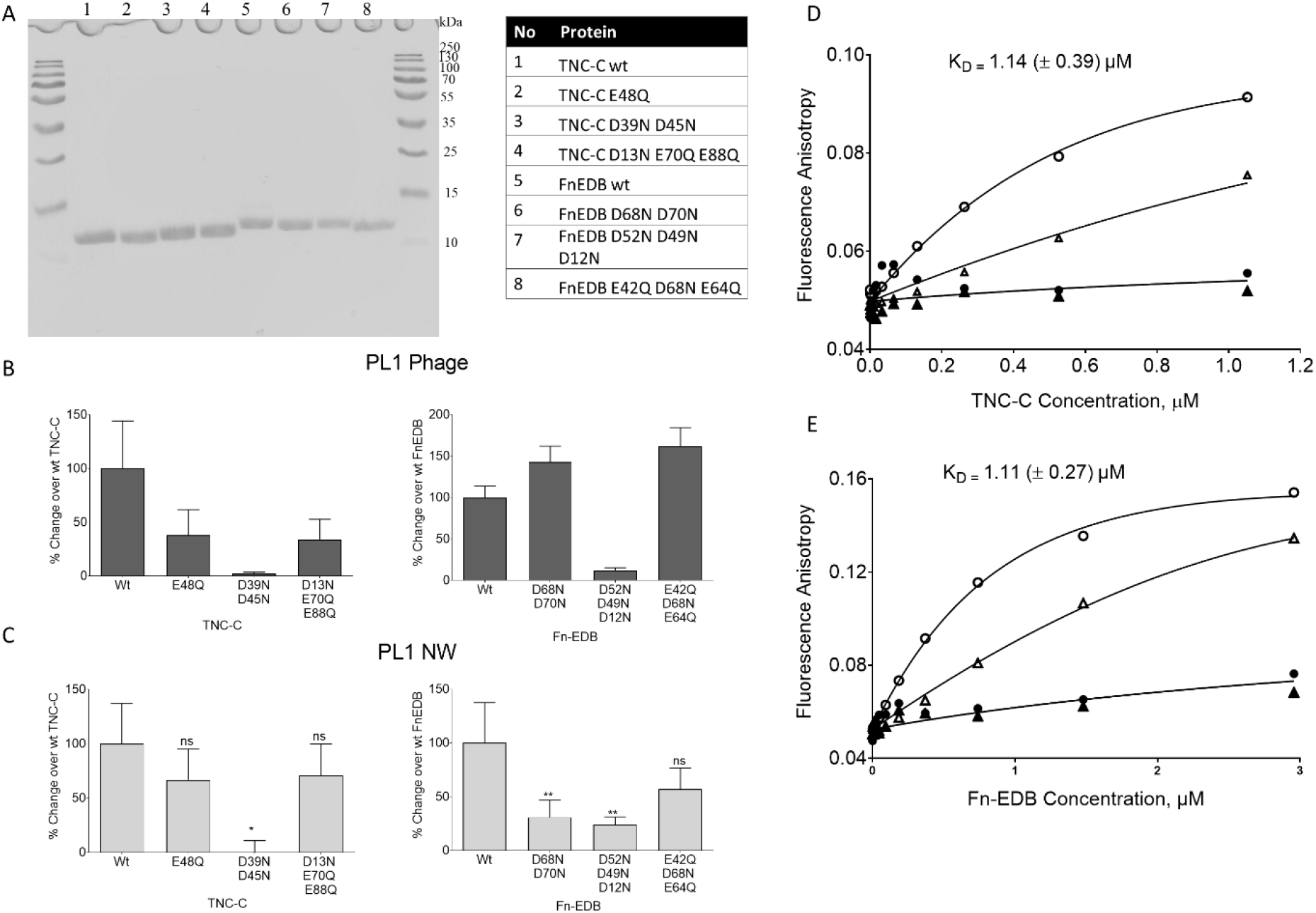
WT and mutant FN-EDB and TNC-C: binding of PL1 peptide NPs and free peptide. (A) The WT and mutant FN-EDB and TNC-C isoforms were expressed in *E. coli.* Coomassie blue-stained SDS PAGE of His-tagged recombinant proteins (loading scheme listed in the table) after purification on HisTrap HP columns. (B) Binding of the PL1 peptide phage to immobilized TNC-C and FN-EDB isoforms. The phage binding is expressed as percent binding over parental WT FN-EDB and TNC-C. (C) Binding of FAM-labeled PL1-NWs to NiNTA beads immobilized recombinant TNC-C and FN-EDB proteins. The mutant TNC-C (D39N, D45N) and FN-EDB (D52N, D49N, D12N) showed a ~ 100% and ~ 76% reduction in binding over WT protein. The PL1-NW binding is expressed as percent binding over parental WT FN-EDB and TNC-C proteins after baseline correction with control protein P32. The binding data are representative of 3 independent experiments. Error bars: mean ± SEM. (D–E) Binding curve characterizing PL1 peptide binding to TNC-C and FN-EDB. FAM-PL1 [0.468 μM (circles) and 4.68 μM (triangles)] was incubated with different concentrations of TNC-C (D) and FN-EDB (E) in the absence (total binding, open symbols) or presence (non-specific binding, filled symbols) of 0.5 mM biotin-PL1. After incubating 24 h at 25 °C, FA values were measured, and binding parameters were calculated as described in Materials and Methods. Data are representative of 3 independent experiments.

We first determined the effect of TNC-C and FN-EDB mutations on engagement with PL1 phage nanoparticles, the biological nanosystem used for biopanning during PL1 identification [3]. PL1-displaying phages showed reduced binding to all TNC-C mutants. In the case of the D39N D45N mutant, binding was reduced by 98%, the effect was milder for the other two mutants (Fig. 3B).

In the case of FN-EDB, the PL1 phage showed a 87% decrease in binding to D52N D49N D12N mutant, whereas PL1 peptide phage binding to FN-EDB D68N D70N and E42Q D68N E64Q mutants was increased by 142% and 161%, respectively (Fig. 3B).

Nanoscaffold can have a profound effect on the targeting ability of the peptide and it is advisable to study the binding using different classes of peptide-guided nanocarriers. Thus, we studied the receptor binding profile of PL1-functionalized iron oxide nanoworms (NWs). NWs were coated with PL1 peptide labeled with FAM fluorophore to enable a fluorescent readout for binding studies. The binding profile of PL1-NWs was similar to PL1-phages, with the strongest reduction of PL1-NW binding to mutant D39N, D45N of TNC-C (~ 100%), and mutant D52N D49N D12N of FN-EDB (~ 76%), respectively (Fig. 3C). These studies mapped the amino acids on FN-EDB and TNC-C involved in engagement with PL1 peptide coated on NPs.

Fluorescent anisotropy allows solution-based characterization of the interaction of small fluorescent ligands with larger binding partners [36] and is thus well-suited for studies of the binding of FAM-PL1 (MW:1.46 kDa) FN-EDB (MW: 12.01 kDa) and TNC-C (MW: 12.3 kDa). The binding of FAM-PL1 to both targets was saturable. The K_D_ values were obtained by global fitting data to a binding isotherm, assuming a single binding site and ligand depletion. For PL1/FN-EDB interaction, the K_D_ was 1.11 ± 0.27 μM (Fig. 3D). For PL1/TNC-C, the fluorescent anisotropy assay and calculations yielded a K_D_ of 1.14 ± 0.39 μM (Fig. 3E). We also observed more extended binding equilibrium kinetic overtime at a decreased concentration of receptors and FAM-PL1 peptide (Fig. S2), suggesting binding slow association kinetics with receptor-ligand interaction sites similar to what was observed *in silico*.

### 2.3. PL1 peptide NPs are internalized in glioma cells via the macropinocytosis pathway

ECM is a dynamic structure and cellular internalization of matrix proteins is increasingly recognized to play a role in ECM remodeling and tissue plasticity [37]. Next, we tested the interaction of cultured U87-MG cells with neutravidin-AgNPs labeled with NHS-CF555 dye and functionalized with biotinylated PL1 peptide. Confocal imaging demonstrated that PL1-AgNPs bound to cultured U87-MG cells, whereas the control biotin-AgNPs showed only background binding comparable to control cells without AgNPs (Fig. 4A-B, S3). PL1-AgNP signal was also present after treatment with a cell-impermeable etching solution to dissolve extracellular AgNPs, suggesting internalization (Fig. 4A). The fluorescence microscopy was supported by flow cytometry data showing that PL1-AgNPs bound and were taken up by U87-MG cells. Control AgNPs showed no significant binding and uptake (Figure 5C-D). Extracellular PL1-AgNPs colocalized with FN-EDB and TNC-C immunoreactivities, whereas no colocalization was seen for the etched cells (Fig. S3B). PL1-AgNP binding and internalization in U87-MG cells was time dependent: we observed only a weak binding at 20 min, whereas progressively stronger binding and internalization were detected after 60- and 180-min incubation with the particles (Fig. S4 A-B).

**Figure 4.**
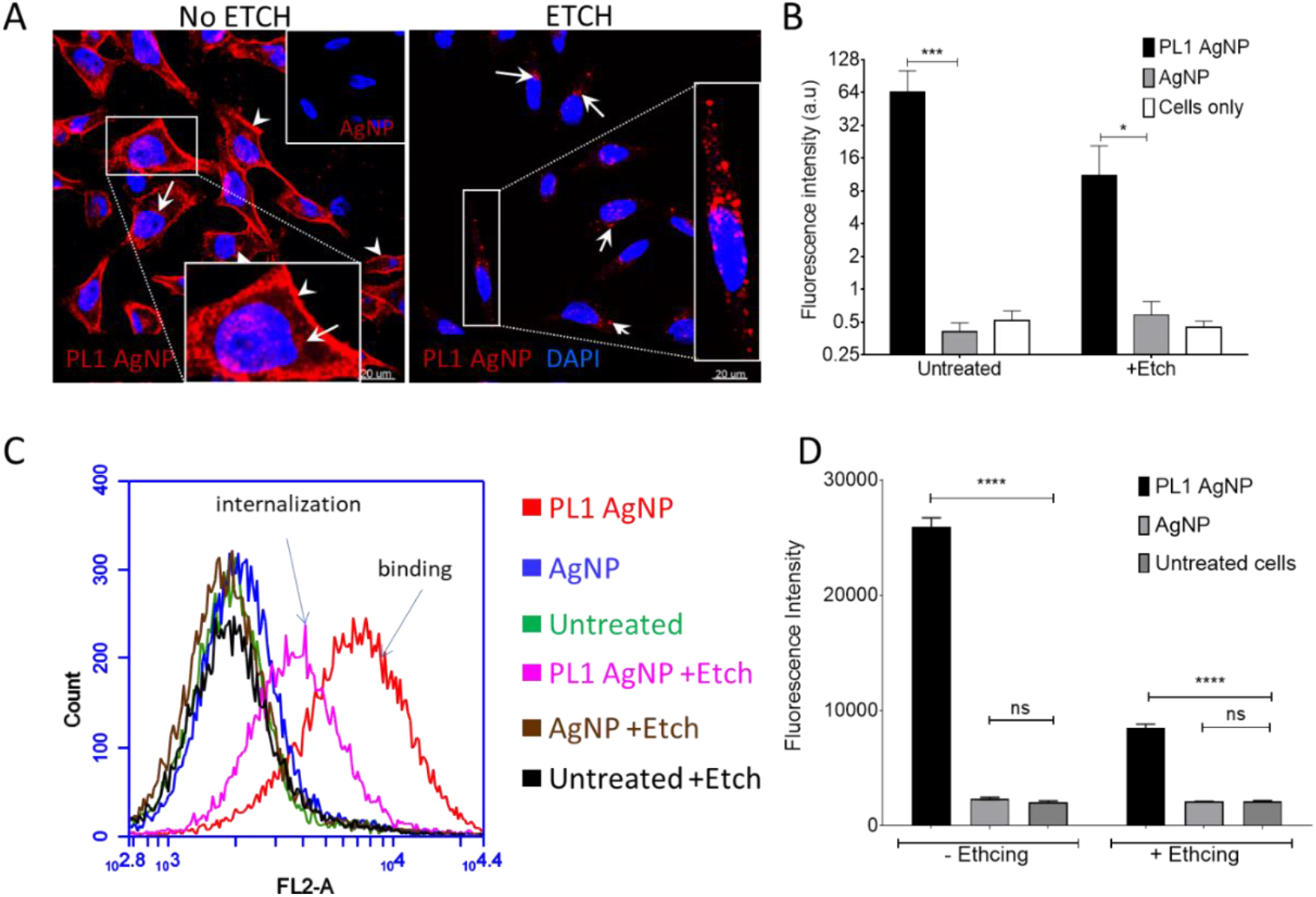
Cellular uptake of PL1-AgNPs. Binding and cellular uptake of PL1 or biotin-blocked CF555-labeled AgNPs in U87-MG cells was assessed by confocal imaging and flow cytometry. (A) Confocal microscopy of U87-MG attached cells incubated with PL1-AgNPs or control AgNPs. U87-MG cells show binding of CF555-labeled PL1-AgNP (red) to the cell surface (white arrowheads), whereas cells incubated with control particles are negative. The etched CF555-labeled PL1-AgNPs show that PL1-targeted particles are able to internalize into the cell cytoplasm (blow-up image shown in box) compare to non-targeted particles. Scale bars: 20 μm (main images) and 2 μm (insets). (B) Binding and internalization into cells was quantified based on CF555 signals in panel (A) of representative images (mean fluorescence intensity). Error bars: mean ± SD (N=12). (C) Flow cytometry of U87-MG cells incubated with AgNPs for 1 h with ± etching. CF555-gated plot showing cell binding and internalization of PL1-AgNP (red: − etch; pink: + etch) compared to control particles AgNP (blue: − etch; brown: + etch). The cells without AgNPs were included as a control standard (green: − etch; block: + etch). (D) Quantification of flow cytometry data were represented as % over control cells without AgNPs, P-value was determined by one-way ANOVA, Error bars: mean ± SEM (N=3), ns P > 0.05, * - P ≤0.05, **** p < 0.0001.

**Figure 5.**
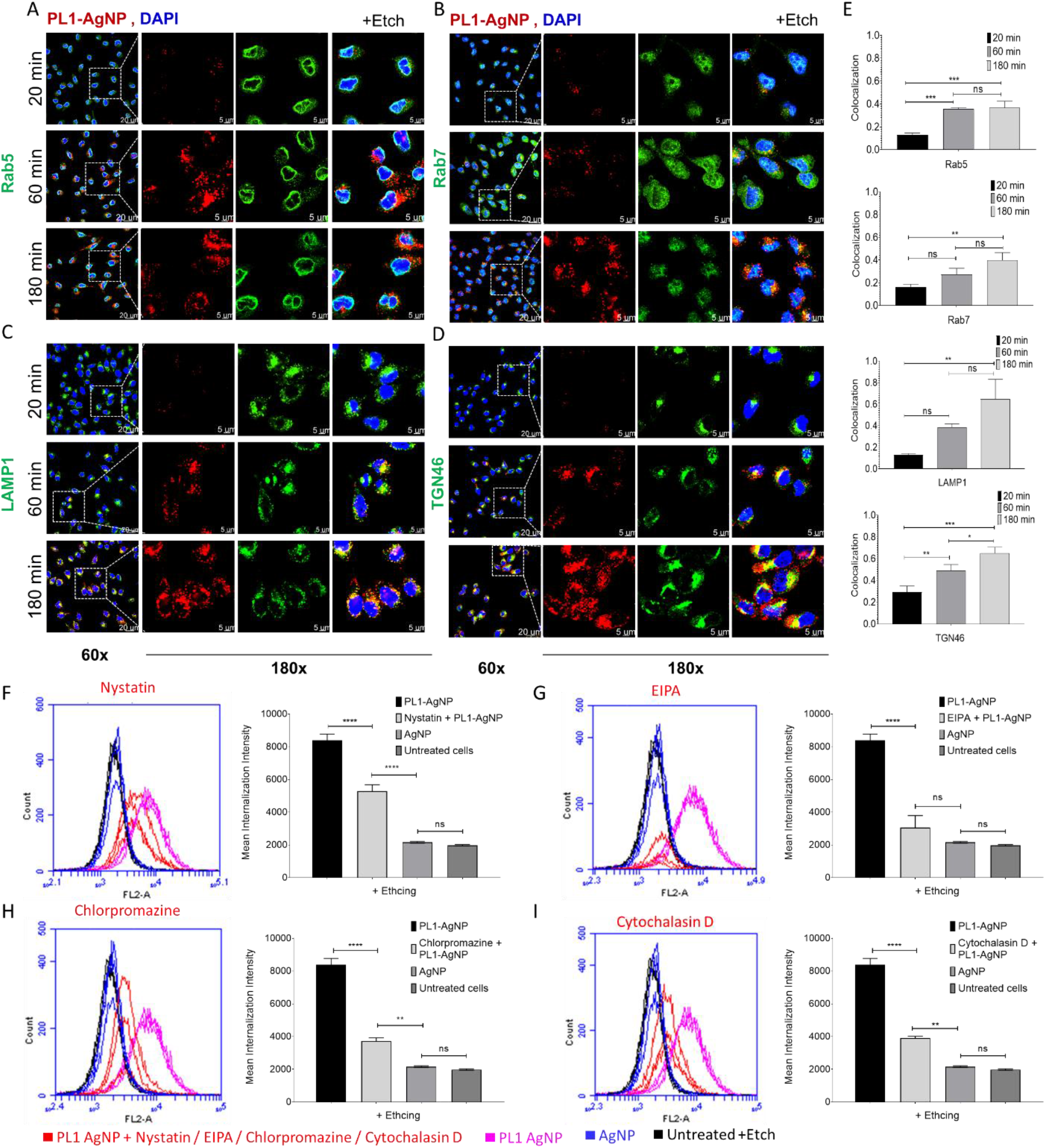
Colocalization of PL1-functionalized silver nanoparticles (AgNPs) with endocytic markers, and inhibition of endocytosis pathways in U87-MG cells. Confocal microscopy images of U87-MG attached cells incubated with PL1-AgNPs (red) at 20, 60 and 180 min time points, and cells were treated by etching solution to visualize only internalized particles. The cells were stained with (A) early endosome marker Rab 5A (green), (B) late endosome marker Rab 7A (green), (C) lysosome marker LAMP-1 (CD107a; green), and (D) trans-Golgi network marker TGN46 (green). The cells were counterstained with DAPI (blue). Scale bar: 20 μm (60x), for the magnified images 5 μm (180x). (E) The relative colocalization was assessed with Manders coefficient (M2) using Image J software and shown in the bar chart. Internalized PL1-AgNP colocalization with markers was quantified from representative images using a One-way ANOVA test. Error bars: mean ± SD (N=3, three independent experiments), ns P > 0.05, *p < 0.05, **p < 0.01, ***p < 0.001, **** p < 0.0001. For the FACS analysis, U87-MG cells were pre-incubated with inhibitors of endocytic pathways, 50 μM Nystatin (F), 1 mM 5-(N-ethyl-N-isopropyl)-Amiloride (EIPA) (G), 10 μM Chlorpromazine (H), 4 μM Cytochalasin D (I), and untreated cells as control, and AgNPs for 30 min. The cells were incubated with PL1-AgNPs for 60 min, and cells were treated by etching to remove membrane-bound particles. The cells were analyzed with flow cytometry for inhibition of uptake. CF555-gated plot showing cell internalization and inhibition of PL1-AgNPs (pink: internalization; red: inhibition) compared to control particles AgNP (blue). The cells without AgNPs were included as a control standard (black). The BD Accuri C6 software was used for the quantification of cell mean intensity from three independent experiments. The bar chart showing one-way ANOVA test, Error bars: mean ± SD (N: 1000 cells/experiment), ns P > 0.05, *p < 0.05, **p < 0.01, ***p < 0.001, **** p < 0.0001.

To map the route of endocytosis of PL1-AgNP, we tracked the colocalization of PL1-AgNPs with compartmental markers: early endosome marker Rab5, late endosome marker Rab7, lysosome marker LAMP-1, and trans-Golgi network marker TGN46. After 20-min incubation, the bulk of PL1-AgNPs was found on the cell surface with very little internalization (Fig. 5A-D, S5 A-D). After 1 h the PL1-AgNPs showed endolysosomal uptake: colocalization with early endosomes positive for RAB5 and some overlap with LAMP-1 and TGN46 (Fig. 5A-D, S5 A-D). At 2 h the PL1-AgNPs showed colocalization with all the tested markers (Fig. 5A-D, S5 A-D), with robust signal also present in trans-Golgi network and lysosomes (Fig. 5B-E, S5 B-D). We next used flow cytometry to study the effect of treatment with endocytic inhibitors (Nystatin, EIPA, Cytochalasin D and Chlorpromazine) on the uptake of PL1-AgNPs in U87-MG cells under conditions established previously [38,39]. We observed that neither Nystatin nor Chlorpromazine nor Cytochalasin D had a significant inhibitory effect on the uptake of PL1-AgNPs, suggesting that PL1-AgNPs are not taken up in U87-MG cells using caveolae/lipid-mediated endocytosis and clathrin-mediated endocytosis pathways (Fig. 5I). In contrast, treatment with two macropinocytosis inhibitors, 5-(N-ethyl-N-isopropyl)-Amiloride (EIPA) and Cytochalasin D, reduced the cellular entry of PL1-AgNPs (Fig. 5I).

## Discussion

Splice isoforms of fibronectins and tenascins, overexpressed in most solid tumors and undetectable in resting adult tissues, have emerged as important targets for affinity ligands for cancer targeting. Recently described dodecameric PL1 peptide specifically engages with FN-EDB and TNC-C overexpressed in solid tumors [3]. PL1 has features that may facilitate translation from preclinical studies to clinical studies: its targets, FN-EDB and TNC-C isoforms, are highly conserved between mice and humans, and show similarly low expression in normal tissues as well as robust upregulation in clinical and experimental solid tumors. Here, we report studies on the characterization of the mode of interaction of PL1 with its receptors and cellular uptake of PL1-guided nanoparticle payloads. In a series of *in silico* modeling and cell-free binding studies we mapped the PL1 target domains in receptor proteins and showed that interaction of PL1 with FN-EDB and TNC-C is driven by a combination of electrostatic interactions and structural features of the peptide. Surprisingly, we observed that PL1-coated NPs were not only deposited on the ECM but internalized in cells via the macropinocytosis pathway.

First, we used a combination of *in silico* modeling and cell-free binding studies to characterize the interaction of PL1 with its receptors. We mapped the sites on receptor proteins that are targeted by the peptide, provided a quantitative estimation of the PL1-receptor binding energies, and determined the nature of the interactions that contribute to the binding. Our data show that the interaction of the peptide with both receptors is primarily electrostatically driven. Molecular dynamics simulation studies suggested that PL1 interacts with acidic residues of both FN-EDB [(D52, D49, D12), (D68, D70), (E42, D68, E64)] and TNC-C [(D13, E70, E88), (D39, D45), (E48)]. Substituting these amino acids with neutrally charged asparagine residues favored unbinding when a harmonic perturbation was applied in order to pull PL1 away from the receptors. PL1 peptide phage or PL1 peptide-NW binding to immobilized recombinant mutant proteins confirmed the critical roles for residues D52N, D49N, D12N in FN-EDB and D39N, D45N in TNC-C. Fluorescence anisotropy-based characterization of interaction of FAM-PL1 with FN-EDB and TNC-C demonstrated stable and saturable binding with a K_D_ of ~ 1.1 μM for both targets – in the same range reported for homing peptides such as RPARPAR [40] and TT1 [41]. Whereas electrostatic interactions are critical for the binding of PL1 peptide to its receptors, structural features of the peptide appear to also be important. A disulfide-bridged version of PL1 peptide with two added flanking cysteine residues has no affinity for either receptor (Lingasamy et al., unpublished). This observation suggest that the flexibility of PL1, observed during the molecular dynamics simulations, is critical for the receptor engagement. Finally, an alanine scanning mutagenesis of PL1 demonstrated important contribution of noncharged amino acids for PL1 engagement with its receptors [3]. Understanding the mode of interaction of PL1 peptide with its receptors developed here serves as the basis for development of improved PL1 derivatives that include nonproteinogenic amino acids with improved *in vivo* stability, enhanced potency, improved tissue distribution, and increased selectivity, as has been reported in the past for other tumor homing peptides [42].

Most anticancer agents exert their cytotoxic effect inside the cells. Therefore, systemic affinity ligands for cancer are typically selected on the basis of their cellular internalization potential. According to a common paradigm, affinity ligands to the ECM components are poorly suited for delivery of intracellularly-acting payloads, necessitating engineering of cleavable or hydrolysable linkers, such as disulfides or hydrazones, for the payload release in the tumor extracellular milieu [43,44]. The confocal imaging and flow cytometry studies on cultured U87-MG glioma cells demonstrated that PL1 peptide NPs not only bind to FN-EDB and TNC-C deposited outside the cells but are also taken up by the cells. PL1-particles were detected at early time points in early endosomes and later in the trans-Golgi network and lysosomes. Mechanistically, the cellular uptake occurred predominantly via macropinocytosis and, to the lower extent, clathrin-mediated endocytosis pathways. The endolysosomal uptake of the PL1-NPs suggests that linker technologies extensively validated with clinical stage internalizing antibody-drug conjugates can be used for coupling anticancer payloads to the PL1 peptide for efficient intracellular release.

The mechanism of cellular uptake of PL1-NPs remains to be determined in detail. Integrins, along with their ECM ligands, are internalized through a variety of endocytic mechanisms, including macropinocytosis [45]. Proteolytic processing of the fibronectin-containing ECM has been shown to facilitate FN endocytosis [46]. It is possible that PL1 hitchhikes on its receptors during cellular internalization at the sites of ECM remodeling. Interestingly and compatibly with simultaneous engagement, PL1 peptide and integrins bind to both FN-EDB and TNC-C at different sites.

Our studies provide basis for optimization of the structure of the PL1 peptide for improved binding and stability as well as provide information for the rational choice of payloads and linker chemistries for PL1-guided therapeutics.

## Conflicts of interest

The data that supports the findings of this study are available from the corresponding author upon request. TT and PL hold a patent on PL1 peptide (“Bi-Specific Extracellular Matrix Binding Peptides and Methods of Use Thereof”; WO. patent no. WO 2020/161602 A1). All other authors declare that they have no competing interests.

## Acknowledgments

We thank the high-performance computing center (HPC) of the University of Tartu and their team for their kind assistance in addressing the technical issues.

## Financial support

This work was supported by the European Regional Development Fund (Project No. 2014-2020.4.01.15-0012), by EMBO Installation grant #2344 (to T. Teesalu), European Research Council grants GBMDDS and GLIOGUIDE from European Regional Development Fund (to T. Teesalu), Wellcome Trust International Fellowship WT095077MA (to T. Teesalu), and Norwegian-Estonian collaboration grant EMP181 (to T. Teesalu). We also acknowledge the support of Estonian Research Council (grant PRG230 and EAG79 to T. Teesalu) and Estonian Research Council grants (IUT20-17 and PSG230).

## SUPPLEMENTARY MATERIALS AND METHODS

**Figure S1.**
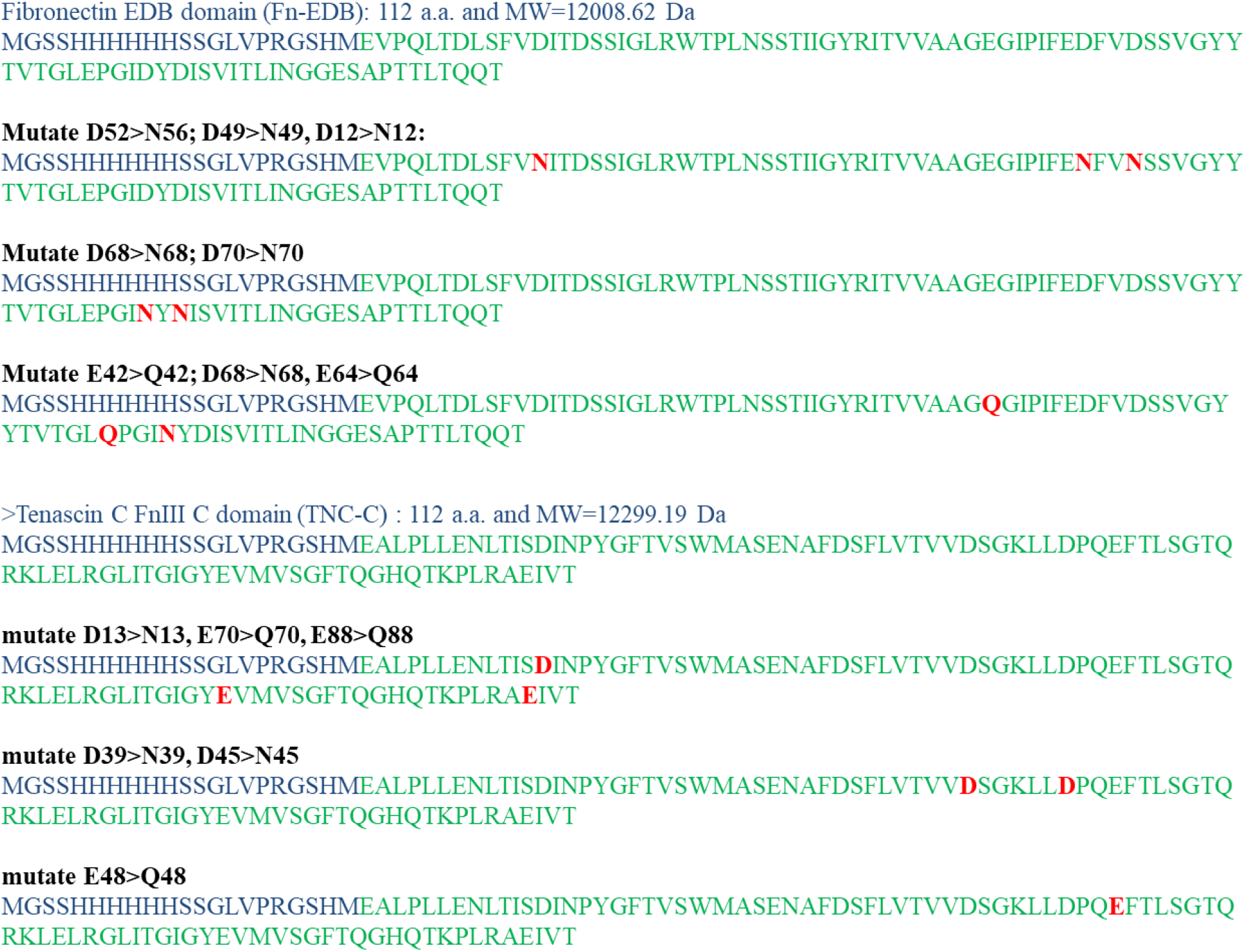
The recombinant FN-EDB and TNC-C WT and mutant protein sequences. The primary sequence of WT proteins (FN-EDB and TNC-C) and respective mutant proteins FnEDB (D68N D70N, D52N D49N D12N, and E42Q D68N E64Q) and TNC-C (E48Q, D39N D45N, and D13N E70Q E88Q). The neutral charge amino acid substitutions are shown in red color.

**Figure S2.**
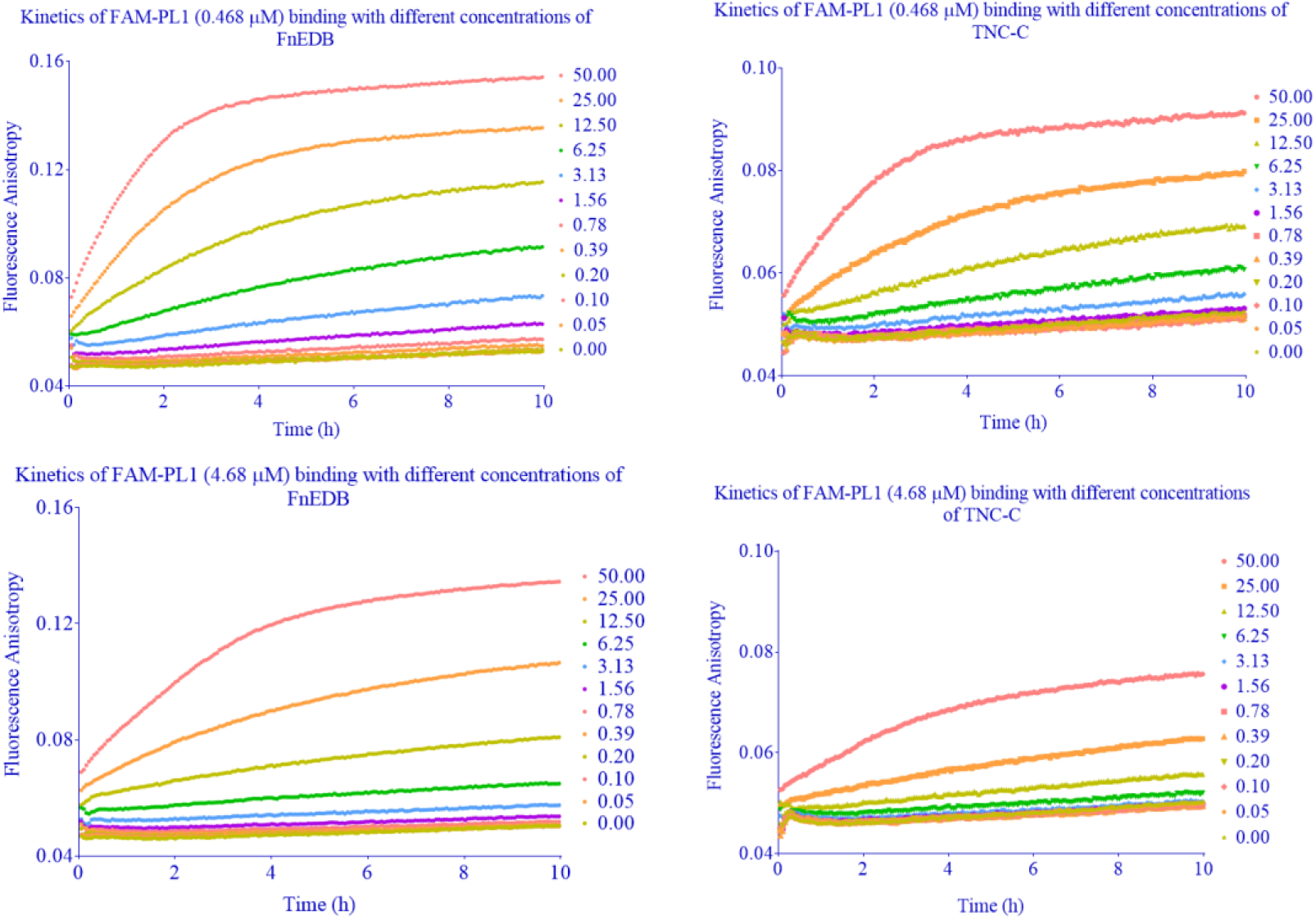
Fluorescence polarization (FP) saturation binding curves. The time course of FA changes caused by FAM-PL1 to different concentrations of FN-EDB and TNC-C. The receptors were prepared from 50 μM to 0 μM per well in the presence of non-specific binding or absence of total binding to 0. 48 μM and 4.68 μM of FAM-labeled PL1 peptide. According to Eq, at the indicated time points, fluorescence intensity was measured, and anisotropy values were calculated. The data representative of experiments from at least three independent experiments.

**Figure S3.**
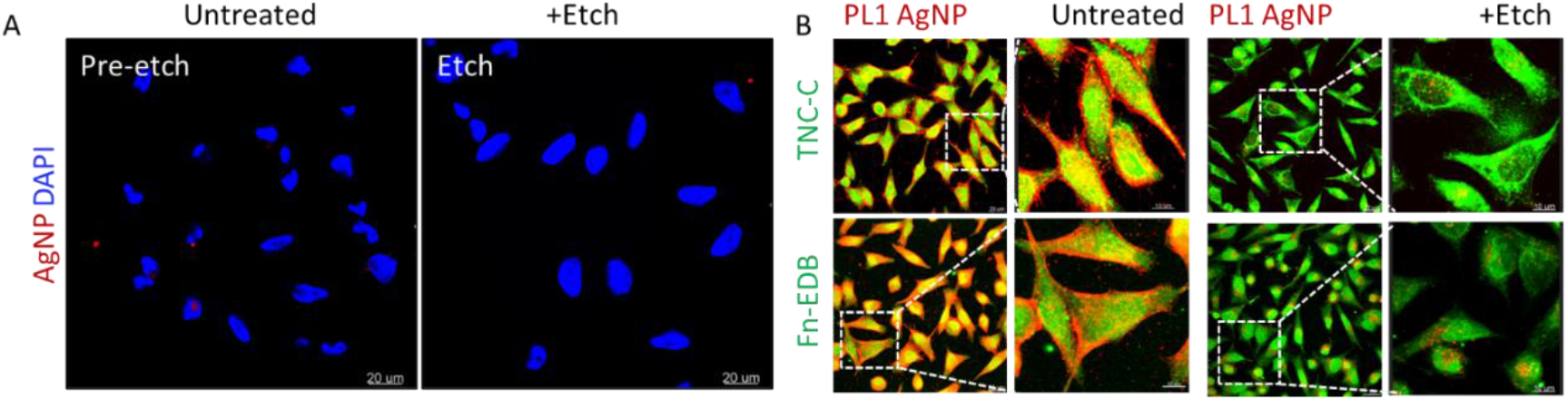
PL1-functionalized silver nanoparticles (AgNPs) colocalize with receptors. (A) Control biotin-AgNPs showed negative binding and internalization in U87-MG cells. Scale bars: 20 μm. (B) The pre-etched PL1-AgNPs (red) colocalize with receptors TNC-C (ScFV G11: Green) and FN-EDB (ScFV L19: Green) Scale bars: 10 μm; magnified images: 5 μm.

**Figure S4.**
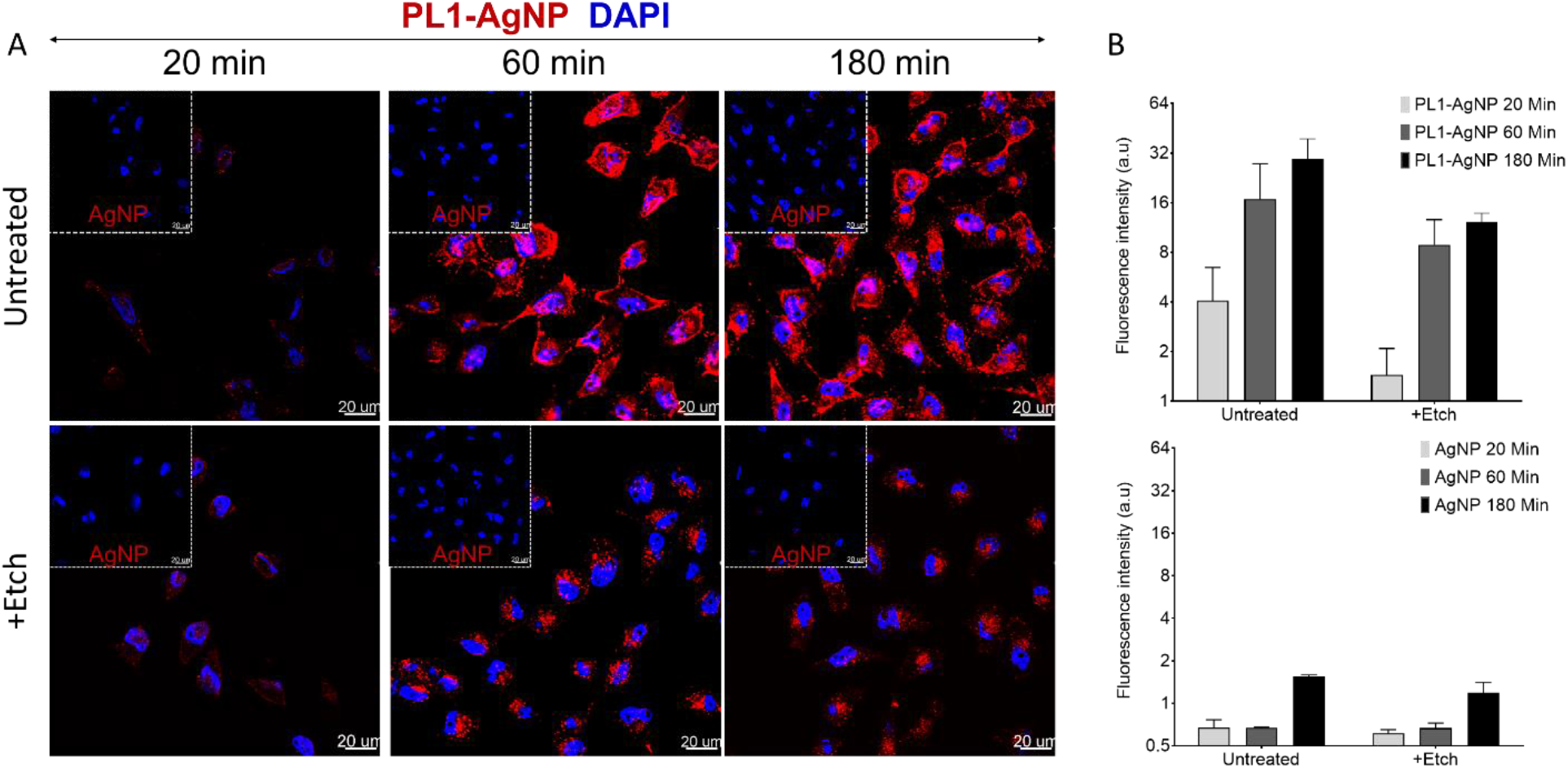
Time-dependent internalization of PL1-functionalized silver nanoparticles (AgNPs) into cells. (A) Confocal microscopy pictures of U87-MG attached cells incubated with PL1-AgNP or AgNP for 20, 60 and 180 min. Some of the cells were etched at 20, 60 and 180 min. The control AgNP samples are completely negative (insets). The pre-etched and etched PL1-AgNPs (red) show superior binding and internalization in a time-dependent manner. (B) Binding and internalization into cells was quantified based on CF555 signal from panel (A) in representative images (mean fluorescence intensity). Error bars: mean ± SD (N=three independent experiments). Scale bars: 20 μm.

**Figure S5.**
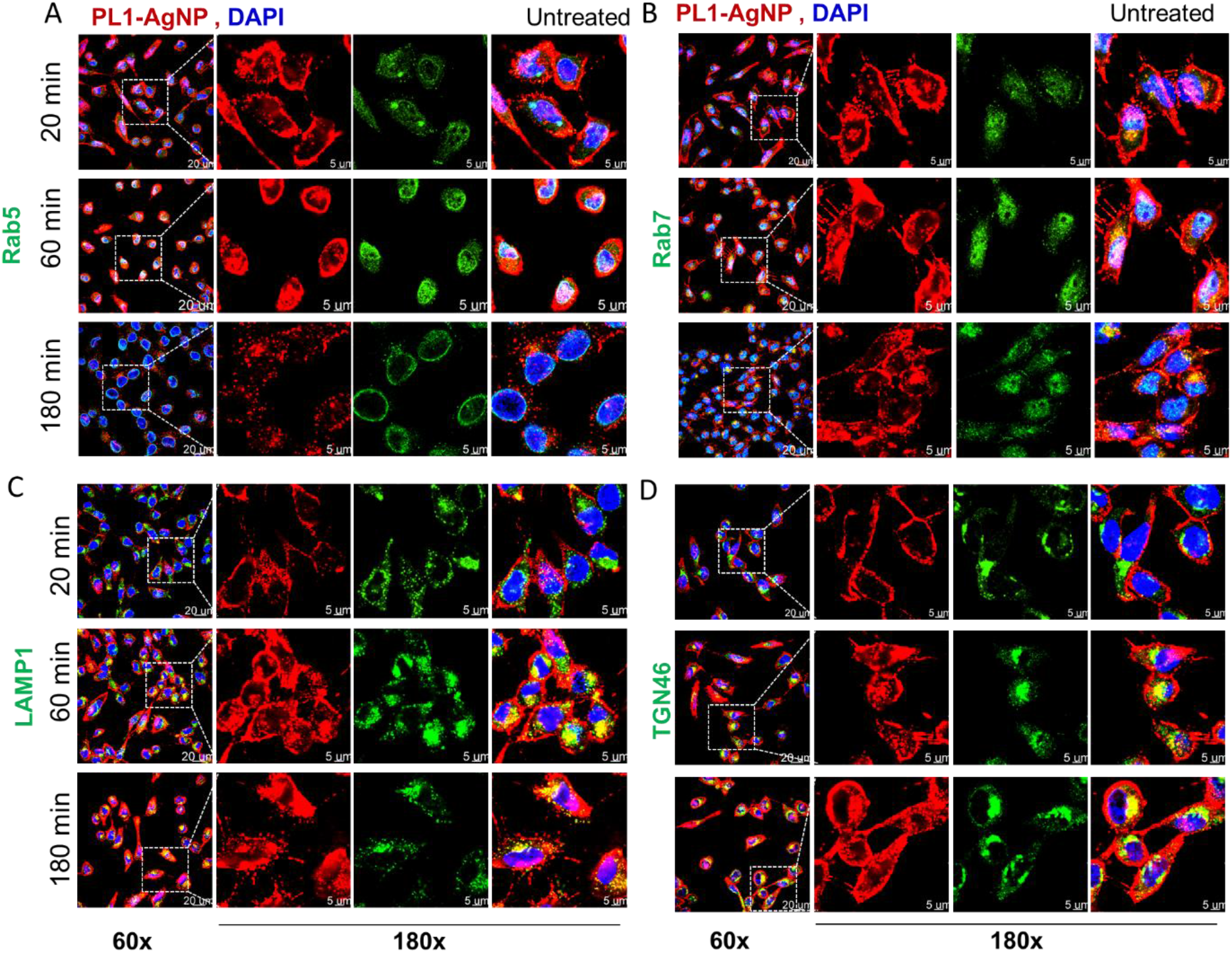
PL1-functionalized silver nanoparticles (AgNPs) colocalization of endocytic markers in U87-MG cells. Confocal microscopy images of U87-MG attached cells incubated with PL1-AgNPs (red) at 20, 60 and 180 min time points. The cells were stained with (A) early endosome marker Rab 5A (green), (B) late endosome marker Rab 7A (green), (C) lysosome marker LAMP-1 (CD107a) (green), and (D) trans-Golgi network marker TGN46 (green). The cells were counterstained with DAPI (blue). Scale bar: 20 μm (60x); magnified images: 5 μm (180x). Representative images from three independent experiments.

## Notes

### Competing Interest Statement

T. Teesalu and P.Lingasamy hold a patent on PL1 peptide (Bi-Specific Extracellular Matrix Binding Peptides and Methods of Use Thereof; WO. patent no. WO 2020/161602 A1).

